# A new leaf sensing organ in a predatory insect group, the praying mantises

**DOI:** 10.1101/2024.04.14.589444

**Authors:** Sydney K. Brannoch, Julian Katzke, Danielle S. Taylor, Evan Economo, Yuri Ogawa, Ajay Narendra, Gavin J. Svenson, Joshua Martin

**Affiliations:** Department of Invertebrate Zoology, Cleveland Museum of Natural History, Cleveland, Ohio, United States of America; Department of Biology, Case Western Reserve University, Cleveland, Ohio, United States of America; Biodiversity and Biocomplexity Unit, Okinawa Institute of Science and Technology Graduate University, 1919-1 Tancha, Onna-son, Okinawa, Japan 904-0495; Department of Biology, Colby College, Waterville, Maine, United States of America; Department of Biological Sciences, Macquarie University, W19F, 205 Culloden Road, Sydney, NSW 2109, Australia

## Abstract

Animals’ sensory systems enable them to navigate and interact with their environments. Adaptive specializations of these systems can generate novel structures or organs that support highly unique niche adaptations. We report the discovery of a novel sensory organ in a group of praying mantises (Insecta, Mantodea, Nanomantoidea), which have an unusual “leaf-planking” ecomorphic life strategy, laying against the undersides of broadleaf vegetation. Histology, scanning electron microscopy, and x-ray computed tomography all support the novelty of this distinct morphology while electrophysiology reveals that the sensory organ, herein designated the gustifolium organ, detects plant volatiles. The location of the gustifolium organon the ventral thoracic surface of these mantises appears to facilitate the chemical detection of the leaves on which it resides. The gustifolium is a novel plant volatile-detecting sensory structure in an obligate predatory insect, directly linked to a newly-identified, highly-adapted life strategy.

## Introduction

Insects possess sophisticated chemosensory capabilities (Hallem *et al*., 2006; Sato & Touhara, 2008, Martin *et al*. 2011, Steele et al. 2023) and much of their behavioral ecology is tied to the detection and discrimination of chemicals (de Boer and Dicke 2006, Xu and Turlings, 2018; Felton and Tumlinson, 2008; Kannan et al. 2022). Perceiving chemical stimuli provides critical information for identifying habitat and oviposition sites, food sources, mates, and predators over short and long distances (Engsontia *et al*., 2008; Iacovone *et al*., 2016). Contact chemoreceptors enable insects to detect the near-field chemical environment via aqueous solutions and dry surfaces (Chapman, 2003), as well as short-range, high-concentration airborne volatiles (Dethier 1972; Stadler 1975; Newland 1998, 1999; Ma et al. 2018). Phytophagous and scavenging holometabolous insects are equipped with such chemosensors (Chapman, 2003), which are typically present as individual receptors on the antennae (Schneider, 1964), mouthparts (Maes, 1977; Hallem *et al*., 2006), wings (Ishimoto & Tanimura, 2004; Hallem *et al*., 2006), tarsi (Hallem *et al*., 2006), and genitalia (Ishimoto & Tanimura, 2004; Hallem *et al*., 2006). The positioning of these receptors is directly related to the insect’s ecological strategy (Zacharuk, 1980). Contact-chemosensory sensilla are located on parts of the body that come into contact, or very near, the source of the chemical signal. For example, the tarsi, and mouthparts of the fly contact food during foraging and a potential mate during courtship, and sensilla that sense nutritients (Ling et al. 2014) or conspecific molecules (Sato and Yamamoto 2020) Selection for highly specialized niches that present unique sensory requirements can lead to the development of novel sensory organs through extensive modification of existing morphology (*e.g*., Vondran *et al*., 1995; Ehlers et al. 2022).

When praying mantis species exploit new habitats, selective pressures for predator avoidance and hunting capability drive the appearance and functionality of the body into specialized, niche-selected morphologies (Svenson & Whiting, 2009; Rivera & Svenson, 2016; Wieland & Svenson, 2018). Species that are similar in morphology, as well as ecological and behavioral strategy, are commonly classified into ecomorphic types (Beuttell & Losos, 1999). Here, we focus on one such ecomorph we are calling “leaf-planking”. These are small, gracile, semi-translucent green praying mantises, which are dorsoventrally compressed, prognathous, and often feature broad, fully developed wings (*e.g*., *Kongobatha* Hebard, 1920, *Enicophlebia* Westwood, 1889) that exhibit an ecological strategy unique within the praying mantises: that adults rest upside down with forelegs held laterad, complanate against the undersides of arboreal broadleaf vegetation (Zhu *et al*., 2002; Tedrow *et al*., 2015; Brannoch & Svenson, 2018),. Furthermore, leaf-planking predation, rest, and flight take-off or landing occurs from the undersides of leaves (S. Brannoch & G. Svenson, personal observation). The leaf-planking ecomorph evolved within a lineage of Old World and Oceanic Nanomantoidea (Brunner de Wattenwyl, 1893; Svenson & Whiting, 2009), comprising two genera of the Leptomantellidae (Schwarz & Roy, 2019) and 20 genera of the Nanomantidae (Brunner de Wattenwyl, 1893; Schwarz & Roy, 2019).

We report the discovery of a new insect chemosensory organ, herein named the “gustifolium organ,” on predatory leaf-planking praying mantises (Insecta, Mantodea, Nanomantoidea). The organ, positioned on the ventral midline of the prothoracic furcasternal plate, contacts the surface of leaves when possessor mantises are at rest. Using histology, electron microscopy, x-ray micro-computed tomography, and electrophysiology (evoked local field potential recordings), we demonstrate that the gustifolium organ is morphologically unique among mantises, responsive to plant-based volatiles, and appears to mediate a novel ecomorphic strategy.

## Results

Leaf-planking mantises adopt a characteristic pose on the ventral surface of a leaf: meso- and metathoracic legs splayed outwards, with the raptorial prothoracic outwards, allowing the body to be pressed close to the leaf surface (Fig. 1A). This foreleg positioning exposes the midline of the furcasternal plate to the surface of the underside of the leaf. In contrast, mantises that perch and hunt on other surfaces adopt different postures (Fig. 1B-D).

**Figure 1.**
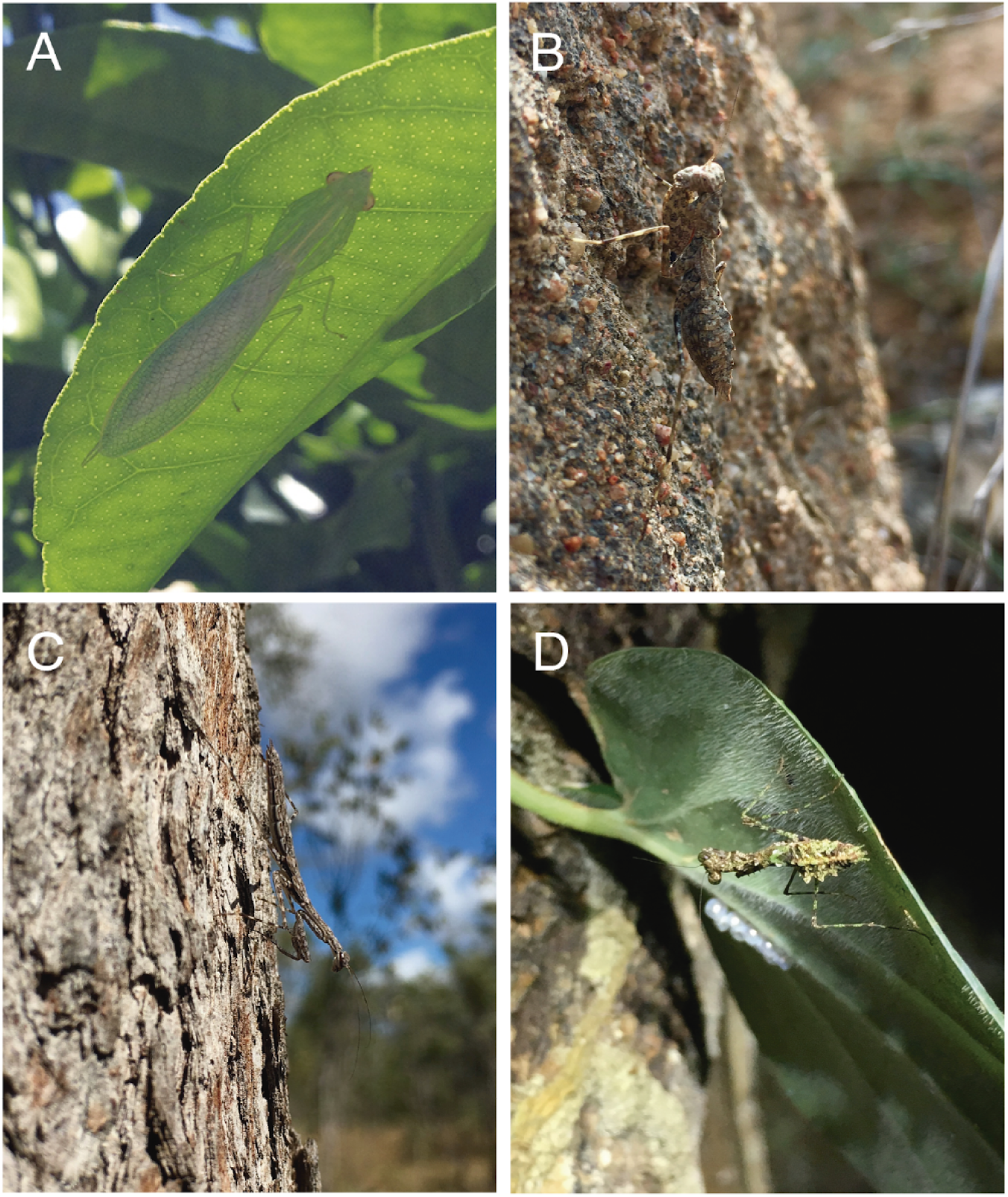
Mantis ecological strategies. *In situ* examples of mantis ecomorph types, (A) *Kongobatha diademata*, leaf-planking. (B) *Phthersigena sp*. ground-dwelling. (C) *Ciulfina sp*. bark-dwelling. (D) *Calofulcinia sp*., generalist perching.

The gustifolium organ is a small, cuticular knob projecting ventrally from the surface of the furcasternum, immediately posterior to the T-shaped sclerite and the insertion sites of the raptorial forelegs on the first thoracic segment (Wieland, 2013; see “furcasternal tubercle” in Brannoch & Svenson, 2016a pg. 74; Brannoch & Svenson, 2016b pg. 16, fig. 20; Brannoch & Svenson, 2018 pg. 6) (Fig. 2). All examined adult and juvenile leaf-planking mantises of both sexes of the study species *Kongobatha diademata* Hebard, 1920, *Leptomantella albella* (Burmeister, 1838), and *Negromantis gracillima* Kaltenbach, 1996, possess this novel gustifolium organ, which is generally sexually monomorphic.

**Figure 2.**
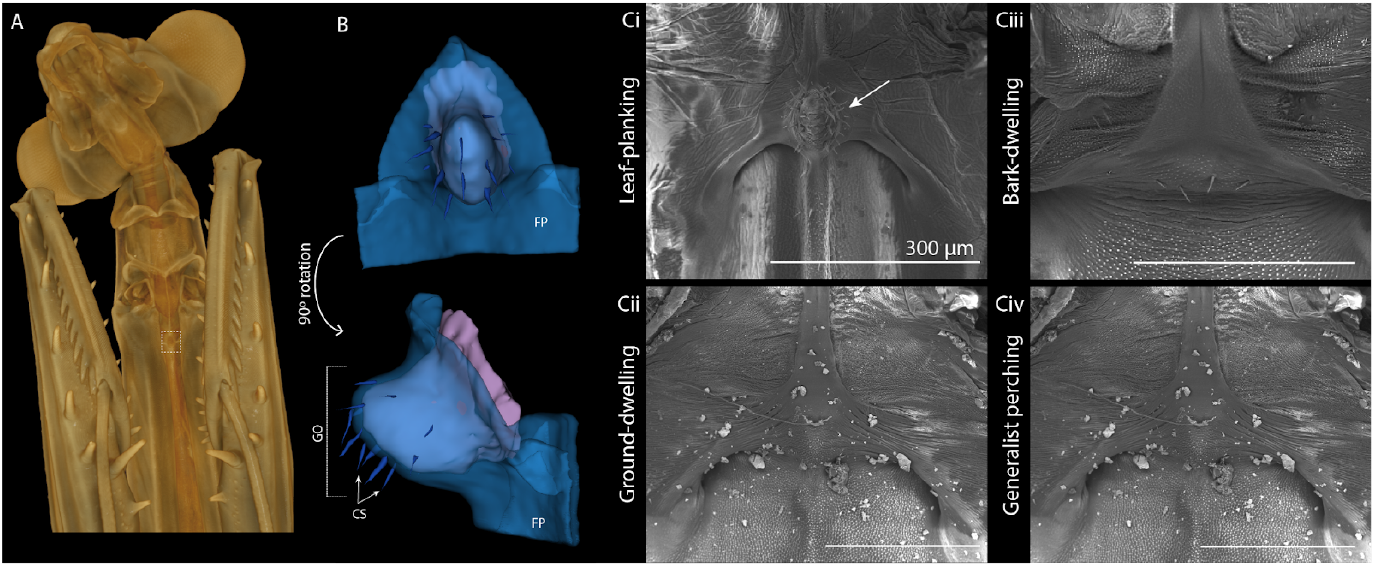
The gustifolium organ. (A) Micro-CT reconstruction of *Kongobatha diademata* (female, ventral view) showing the location of the gustifolium organ between the forecoxae (white dotted box). (B) Segmentation of the gustifolium organ with cuticle (transparent blue), chaetic sensilla (royal blue), and neural tissue (purple); ventral view (anterior segmentation), lateral view (posterior segmentation). (C) Scanning electron micrographs of (i) leaf-planking *Kongobatha diademata*, female, (ii) ground-dwelling *Paraoxypilus sp*., female, (iii) bark-dwelling *Amorphoscelis pulchella*, female. **Abbreviations** CS = chaetic sensilla, FP = furcasternal plate, GO = gustifolium organ

All other species, surveyed across several extant Mantodea families (examples in Fig. 2C i-iv), had either the common T-shaped sclerite structure found in all praying mantises (Wieland, 2013), or a much less pronounced structure with a few sensilla (e.g. Fig 2C iii). The gustifolium organ proper was limited to species of leaf-planking mantises.

Approximately 14 to 25 sensilla surface the organ, which curve slightly and are grooved, tapered apically, and feature a tip pore (Fig. 3a–c). These sensilla are situated in flexible, membranous sockets (Fig. 3c) that would allow mechanical deflection to occur, suggesting that they may also serve a mechanoreceptive function. The structural characteristics of the organ’s sensilla correspond to chaetic sensilla present on the antennae of mantises and other insects (Zacharuk, 1985; Chapman, 1998; Carle *et al*., 2014). Chaetic sensilla are typically contact chemoreceptors, mechanoreceptors, or a combination of those senses (Zacharuk, 1985; Carle *et al*., 2014).

**Figure 3.**
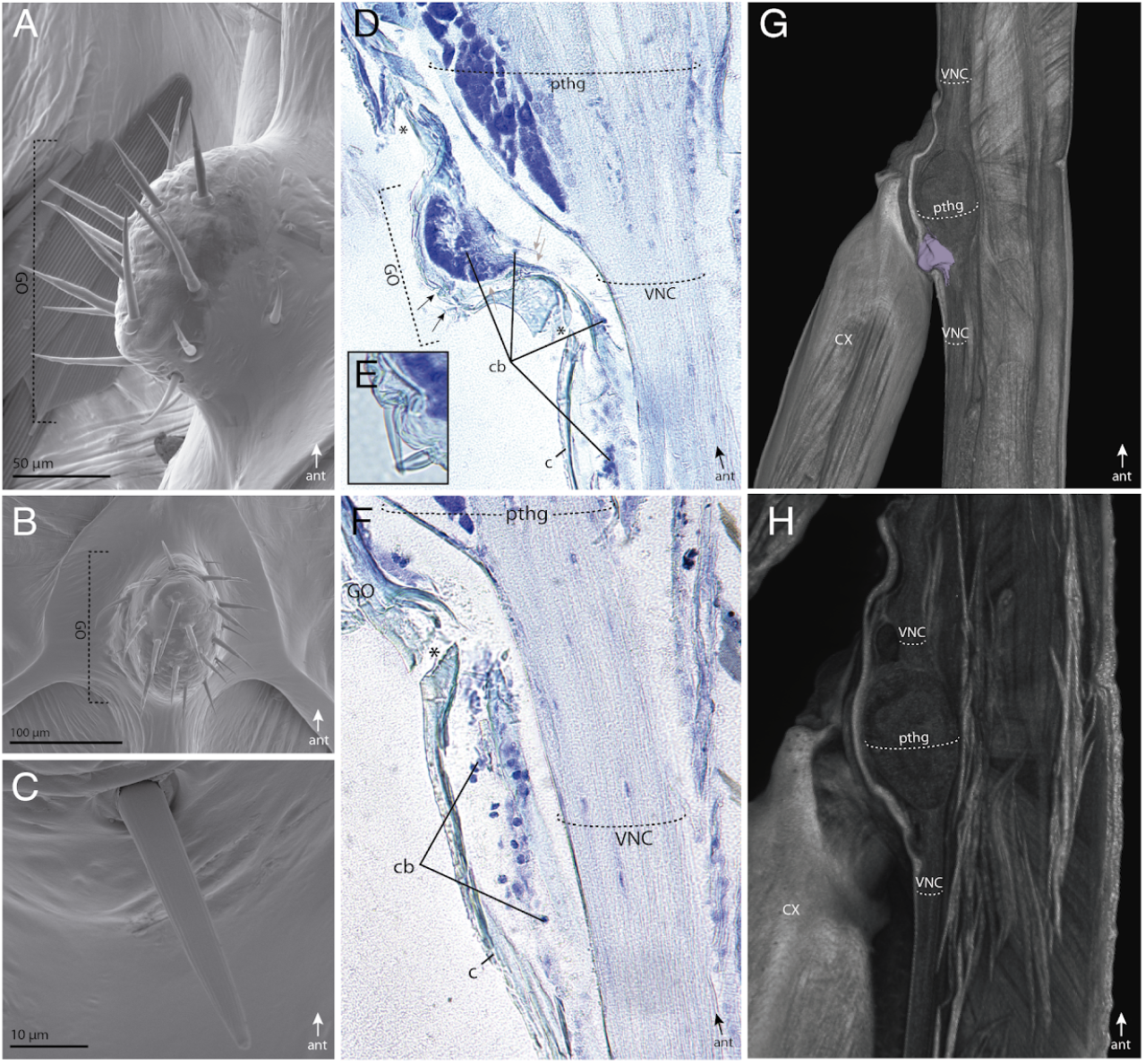
External and internal morphology of the gustifolium organ (GO). (A-C) Scanning electron micrographs (SEM) of *Kongobatha diademata* (male) GO: (A) lateral view, (B) ventral view, (C) close up of a sensillum. (D-F) Midsagittal histological section (12µm) of *Leptomantella albella* (female) GO stained with toluidine blue. Dark blue regions are the somata of neurons; chaetic sensilla (black arrows), nerve fibers (gray arrows) and cuticle broken during the sectioning process (asterisks) are labeled on the image. (G) Midsagittal micro-CT volume rendering of leak-planking *Negromantis gracillima* prothorax with neural tissue of the GO (purple). (H) Midsagittal micro-CT volume rendering of plant-associated *Orthodera sp*. prothorax. **Abbreviations:** ant = anterior; c = cuticle; cb = cell bodies; CX = forecoxa; GO = gustifolium organ; pthg = prothoracic ganglion; VNC = ventral nerve cord.

The internal structure of the organ and surrounding region were captured using histological sectioning and X-ray micro-computed tomography (micro-CT), which illustrates the ventrally projected, pocketed outgrowth along the midline of the furcasternum that forms the cuticular structure of the organ (Figs. 2b, 3d, 3g). The internal cavity of the gustifolium organ is filled with neuronal cell bodies with axons projecting toward the ventral nerve cord (Fig. 3d–f). A gap between the ventral body wall of the furcasternum and the prothoracic ganglion was observed in leaf-planking *K. diademata* (Fig. 3d–f). In addition, morphological data from a plant-associated mantis from a non-leaf planking family (*Orthodera* Burmeister, 1838) points to a potential ancestral morphological state for Mantodea with the prothoracic ganglion abutting the ventral body wall with a minimal gap, and a furcasternum without the pocketed outgrowth found in the gustifolium (Fig. 3h).

Local field potential recordings from the neurons within the gustifolium organ of intact, leaf-planking female *Kongobatha diademata* specimens were conducted to examine response characteristics to external stimuli to confirm 1) if the organ is chemosensory, and 2) its specific sensitivities to relevant compounds (Fig. 4). Plant volatiles, hexane, and untreated air were delivered as a laminar air stream across the sensilla of the gustifolium organ. Voltage changes in response to the onset of a stream of odor-free air (control) were not different from zero (Fig. 4Bi, C; one-sample Wilcoxon signed rank test p<0.05), confirming that the organ did not respond to the mechanical stimulation from the air flow.

**Figure 4.**
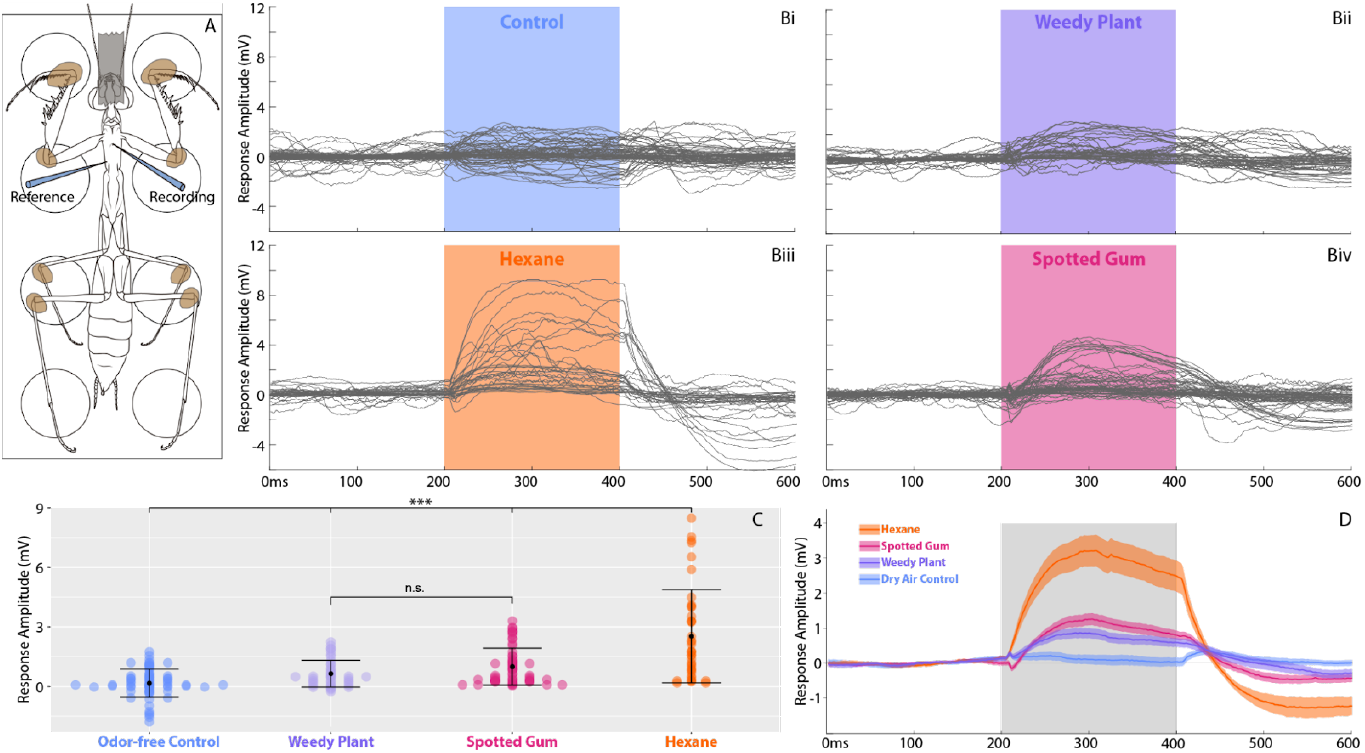
Electrophysiological responses to odor stimulation. (A) Extracellular recording schematic: specimens were affixed to a block with beeswax, and the head capsule was affixed with tape. A glass electrode (Recording) was inserted into the gustifolium organ, and a reference electrode (Reference) was inserted into the prothorax a few millimeters away. From the base of the recording set-up, the stimulus air flow was directed anteriorly. The antennae were removed before stimulation began. (B) Evoked local field responses to (i) clean, odor-free air control (blue), (ii) weedy plant leaf volatiles (purple), (iii) hexane (orange), and (iv) spotted gum leaf volatiles (pink). The shaded, colored area indicates the stimulus period. All individual responses are shown (seven specimens, five trials for each stimulus). (C) Peak response amplitude for all specimens, all trials, for each of the four stimuli. Responses to all stimuli were different from control, and responses to hexane were different from responses to weedy plant and spotted gum volatiles (^***^, p <0.005, Tukey HSD). (D) Average responses (line) +/-the standard error of the mean (shaded) across all specimens and trials.

*K. diademata* has been observed perching on the leaves of spotted gum (*Corymbia maculata*) (S. Brannoch, personal observation). In response to stimulation with volatiles from spotted gum leaves, the local field potential increased for the duration of the stimulus, then dropped below the baseline when stimulation ended (Fig. 4Biv, D). The mean response to volatiles from a weedy ground plant (common ground epiphyte) was similar, with a slightly lower amplitude (Fig. 4 Bii, D). Stimulation with hexane produced the largest response, with a similar increase above baseline during stimulation, and decrease below baseline after the stimulus ended (Fig. 4 Biii, D). The mean response to hexane, spotted gum, and weedy plant volatiles were all different from control, and hexane was different from both spotted gum and weedy plant volatiles (p<0.005, Tukey multiple comparisons).

## Conclusions

The gustifolium organ is a novel chemosensory organ found in mantises who occupy a novel leaf-planking ecomorph. We have described the unique anatomy, location, and connection to the thoracic ganglia of this organ, which is not found in similar mantises. The organ is covered in chaetic sensilla, which play a role in contact chemoreception and mechanosensation in other insects (Hallberg and Hansson 1999). The concentrated, long-duration stimulation with volatiles mimics the close contact the organ would have with a leaf. Similar responses to higher concentrations of air-born volatiles have been reported in contact chemoreceptor sensilla in other insects (e.g. the blowfly, Chapman, 2003). The lack of response to untreated air flow, which would ostensibly deflect the chaetic sensilla and elicit a neural response in mechanoreceptor neurons, suggests that the gustifolium organ may not respond to mechanical stimuli. However, the location of the sensilla suggests that they could monitor contact of the ventral surface of their body with a substrate, and perform a dual role. We conclude that the gustifolium is a dedicated, novel sensory apparatus for detecting leaf volatiles in leaf-planking mantises.

We have named the gustifolium organ for its putative leaf “tasting” capabilities. We hypothesize that the organ facilitates substrate determination in leaf-planking praying mantises. As leaf-plankers exhibit a unique ecomorphic strategy relative to other praying mantises, in that they reside flattened against the undersides of tree leaves, this substrate determination is tuned to specific plant-based chemical information associated with their leaf dwelling lifestyle. This function is a profound departure from traditional notions of praying mantises living a free-roaming, predatory lifestyle. It suggests that insect predators can orient not only to a specific habitat, but to a particular perch on a host plant, using sensory input beyond visual cues and olfactory sensilla on the antennae. While the evolution of specialized sensory organs for plant host detection has been documented for phytophagous insects (*e.g*., infrared receptors in pyrophilic Buprestid beetles, Vondran *et al*., 1995), the phenomenon is not only rare, but to our knowledge has never been documented in predatory insects. The existence of the gustifolium organ raises the question as to why a mantis lineage evolved an entirely new structure rather than utilize or expand on existing sensory morphology that could provide the same sensory cues. We suggest that the ecology of the leaf-planking lineage was so unique in the history of mantises that it required its own, dedicated sensory channel.

## Materials and Methods

### Scanning Electron Microscopy (SEM)

Dry preserved, unpinned *Kongobatha diademata* were sputter coated with palladium and imaged with a FEI Nova Nanolab 200 Field Emission SEM and a FEI Helios Nanolab 650 SEM. Images were processed in Adobe Photoshop CC 2015.5 to adjust brightness and contrast and were plated in Adobe Illustrator CC 2015.3.

*Histology*. An ethanol (95%) preserved specimen of the leaf-planking species *Leptomantella albella* was dissected to isolate the prosternum. The specimen was rehydrated in 90% and 80% ethanol for four hours each before being placed in 70% ethanol for 2 days. The specimen was then fixed in acetic acid-alcohol-formaldehyde solution for three days. The specimen was dehydrated in graded ethanol treatments of fresh 70%, 90%, and 95% for 15 minute intervals, followed with three thirty minute intervals of fresh 100% ethanol. The specimen was submerged in a 1:1 solution of 20 ml Xylene and 100% ethanol for 30 minutes and was then submerged in xylene solution for two thirty minute intervals, the last 20 minutes of which took place in a 60ºC oven. The specimen was then wax embedded and serially sectioned using a Reichert Ultracut microtome with glass knives, resulting in 12 μm thick sections. Slides were stained with 1% Toluidine Blue, which stains neural tissue dark blue (Feirabend *et al*., 1998; Thompson & Roosevelt, 1998). Slides were cover-slipped with a drop of Cytoseal 60.

### X-ray Micro-Computed Tomography

Micro-CT scans were conducted with a ZEISS Xradia 510 Versa 3D X-ray microscope with accompanying ZEISS Scout and Scan Control System software (version 10.7.2936), based on 1600 increments over 360 degree rotations, at the Okinawa Institute of Science and Technology Graduate University, Japan. Scanned specimens included 95% ethanol preserved material accessioned within the Cleveland Museum of Natural History (see Table 1). Specimens were stained with 4% Iodine (I_2_, 40 mg/ml) solution for at least 24 hours or a 1% PTA (phosphotungstic acid, 10 mg/ml) solution in 99% ethanol for at least seven days. Immediately prior to scanning, specimens were transferred to small tubes filled with 99% ethanol. To yield the highest scan quality, scan settings were specifically chosen for each specimen. 3D reconstructions were performed with Zeiss Scout-and-Scan Control System Reconstructor software (version 11.1.6411.17883). Output files were saved in DICOM format.

### 3D Analysis

The micro-CT scans resulted in image stacks that were processed and examined using 3D Slicer (https://www.slicer.org/, accessed: 2018 April 29; Fedorov *et al*., 2012; Kikinis *et al*., 2014) in three two-dimensional planes (*i.e*., axial (x-y), sagittal (y-z), and coronal (x-z)). Cuticle, sensilla, and neural tissue comprising the gustifolium organ were manually segmented using the Segment Editor Module in order to create three-dimensional (3D) reconstructions. A transparent rendering of the prothorax was created using the Volume Rendering Module, with the display preset CT-MIP and VTK GPU Ray Casting setting. Scalar color was mapped according to the black, gray, and white color intensity values for each individual scan, which enabled for the adjustment of the scalar opacity mapping according to those set color values. Further volume renderings were created with Drishti v2.6.4 (Limaye, 2012). Structures were masked in Amira 6.5 (Thermo Fisher Scientific) and then imported as individual volumes into Drishti v.2.6.4. Resultant figures were plated in Adobe Illustrator CC 2015.3.

### Electrophysiology. Kongobatha diademata

Hebard, 1920 individuals were collected by net and light trapping techniques in Kuranda and Daintree Rainforest Observatory, Queensland, Australia June–July 2016, February 2017, and January 2018 (see Brannoch *et al*. 2017 for description of praying mantis collecting techniques). Live mantises were individually housed in a climate-controlled room at 25°C (77°F) with 63% humidity in plastic containers with fine mesh tops for air flow. Mantises were misted with water daily, fed cultured *Drosophila* 3–4 times a week, and were kept at a 12:12 hour light-dark schedule using ∼300–650 nanometer halogen metal halides.

Live *Kongobatha diademata* individuals were cold anesthetized prior to being tethered ventral side up to a Lego brick. To ensure immobility of the specimen throughout the experimental procedure, specimens were fixed to the brick’s studs at the prothorax, abdomen, and the femoral-tibial and tibial-tarsal joints of the legs via low melting point beeswax while the head capsule was fixed with a thin piece of tape. The antennae were excised at the base of the scape to prevent stimulation of antennal olfactory receptors. A fine glass electrode was filled with phosphate-buffered saline and the tip was coated with an electrically conductive gel (Spectra ® 360 electrode gel). The electrode was inserted into the cuticle of the gustifolium organ with a glass reference electrode inserted in the cuticle of the base of the furcasternal plate, both of which were held in place using micromanipulators (see schematic in Fig. 4). The electrophysiology setup was stabilized using a Kinetic Systems 8001 stabilizing breadboard table to minimize vibrations. Extracellular activity of the gustifolium organ was recorded *in vivo* under four experimental conditions: laminar air flow, hexane treated laminar air flow, *Corymbia maculata* treated laminar air flow, and weedy ground plant treated laminar air flow.

Stimulus presentations were controlled with a Syntech stimulus controller CS-55 system (The Netherlands), which controlled air flow rate and duration. Tubing connected to the Syntech stimulus controller CS-55. For the control, untreated air was pumped through a glass capillary. For the hexane treatment, air was pumped through a glass capillary within which a piece of filter paper freshly saturated with Hexane (Sigma-Aldrich PTY. LTD. (AUS)) was placed. For the *Corymbia maculata* treatment, air was pumped through a glass capillary within which a crumpled *Corymbia maculata* leaf was placed. For the *weedy plant treatment*, air was pumped through a glass capillary within which a crumpled leaf from a common ground epiphyte was placed. Controlled puffs of air with consistent airflow force and duration were produced by a Syntech stimulus controller CS-55. The flow rate of each air puff was measured with a Cole-Parmer air flow meter, totaling 1.8 liters per minute.

Using 4 female *Kongobatha diademata* specimens, a total of 5 stimulus presentations took place for each stimulus type (air flow, hexane treated air flow, *C. maculata* treated air flow, and weedy ground plant treated air flow), followed by a 5 minute rest period. Each stimulus presentation was 200 milliseconds in duration with an interstimulus period of 10 seconds, totaling 1 minute of recording time. The order of each stimulus type was randomized. Physiological recordings were performed with a Syntech IDAC4 system. Scripts were written in MATLAB 2014a.

## Acknowledgements

The authors thank Mariella Herberstein, Kate Barry, and Nathan Hart of Macquarie University and Al Pollack of Case Western Reserve University for aid in preliminary electrophysiological experiments. The authors thank Polychronis Rempoulakis and Md. Jamil Hossain Biswas of Macquarie University and George Fischer and Francisco Hita Garcia of Okinawa Institute of Science and Technology (OIST) for their training on equipment necessary for the completion of this project. The OIST Imaging Section provided access to the micro-CT scanner. The authors thank Michele Schiffer of Daintree Rainforest Observatory (James Cook University, Queensland, Australia); Zoe-Joy Newby of the Australian Botanic Garden; Nanthawan Avishai and Danqi Wang of the Swagelok Center for Surface Analysis of Materials, Case Western Reserve University. One author (SKB) would like to thank David Rentz for his assistance with field work in Queensland, Australia. This project was supported by the National Science Foundation East Asia and Pacific Summer Institute (EAPSI) under the grant OISE-1614151, the Student Research Travel Award from the Systematics and Evolutionary Biology section of the Entomological Society of America, the Phi Beta Kappa Research Grant (Case Western Reserve University) awarded to Sydney K. Brannoch, and subsidy funding to OIST. This project was further supported under the NSF grant DEB-1216309 to Gavin J. Svenson and the NSF grant IIS-1704366 to Joshua P. Martin. Any opinions, findings, and conclusions or recommendations expressed in this material are those of the authors and do not necessarily reflect the views of the National Science Foundation, the Entomological Society of America, or Phi Beta Kappa.

## Author Contributions

S.K.B. discovered the gustifolium organ, collected and analyzed morphological data, performed extracellular recordings, and wrote the manuscript; J.K. performed micro-computed tomography scans and contributed in writing; D.S.T. assisted in analyzing electrophysiological data and in writing the manuscript; E.E. contributed intellectually and in writing; Y.O.K. assisted in project design and extracellular recordings; A.N. assisted in project design; G.J.S. contributed intellectually and to the writing of the manuscript; J.P.M. assisted in project design, analyzed electrophysiological data, and wrote the manuscript.

## Notes

### Competing Interest Statement

The authors have declared no competing interest.

